# Lymphoid follicle formation and human vaccination responses recapitulated in an organ-on-a-chip

**DOI:** 10.1101/806505

**Authors:** G. Goyal, P. Prabhala, G. Mahajan, B. Bausk, T. Gilboa, L. Xie, Y. Zhai, R. Lazarovits, A. Mansour, Min Sun Kim, D. Curran, J. M. Long, S. Sharma, L. Cohen, O. Levy, R. Prantil-Baun, D.R. Walt, D.E. Ingber

## Abstract

Lymphoid follicles (LFs) are responsible for generation of adaptive immune responses in secondary lymphoid organs and form ectopically during chronic inflammation. A human model of LF formation would provide a tool to understand LF development and an alternative to non-human primate models for preclinical evaluation of vaccines. Here, we show that primary human blood B- and T-lymphocytes autonomously assemble into ectopic LFs when cultured in a three-dimensional (3D) extracellular matrix gel within an organ-on-a-chip microfluidic device. Dynamic fluid flow is required for LF formation and prevention of lymphocyte autoactivation. These germinal center-like LFs contain B cells expressing Activation-Induced Cytidine Deaminase and exhibit plasma cell (PC) differentiation upon activation. To explore their utility for vaccine testing, autologous monocyte-derived dendritic cells were integrated into LF Chips. The human LF chips demonstrated improved antibody responses to split virion influenza vaccination compared to 2D cultures, which were enhanced by addition of a squalene-in-water emulsion adjuvant, and this was accompanied by increases in LF size and number. When inoculated with commercial influenza vaccine, PC formation and production of anti-hemagglutinin IgG were observed, as well as secretion of cytokines similar to those observed in vaccinated humans over clinically relevant timescales.

## INTRODUCTION

New vaccines and immunotherapies are currently evaluated in animal models, which can lead to unpredicted toxicities or poor efficacy in clinical trials because of species-specific differences in immune responses^1^. Preclinical experiments can be conducted *in vitro* using human immune cells collected from blood; however, even these results often fail to predict patient responses^2, 3^. One major reason for this failure is that *in vivo* immune responses commonly occur within the highly specialized tissue microenvironment of lymphoid follicles (LFs) that normally exist within the cortex of secondary lymphoid organs such as lymph nodes. However, they also can form ectopically (in other organs) as a result of inflammation where they are similarly responsible for generating an adaptive immune response^4^. Importantly, DNA modifying enzymes required by B cells to switch from one Ab isotype to another, such as Activation-Induced Cytidine Deaminase (AID), are only expressed by B cells resident within LFs^5^. Upon immune activation by antigen, helper T cells in the LF enable the B cells to switch isotypes and produce high affinity antibodies (Abs); when activated in this manner, LFs are called germinal centers^6^. The importance of this three-dimensional (3D) tissue niche is made clear by the observation that class switching is defective in mice or humans that lack lymph nodes, and this leads to recurrent infections and morbidity even though isolated B-lymphocytes from these animals or patients can still be induced to undergo class switching in vitro^7^.

Thus, creation of a well-defined experimental model that can recapitulate human LF formation *in vitro* could provide insight into how these structures that are so critical for immune responses develop. Given the current challenges related to the availability of animal models, and particularly non-human primates, for testing vaccines for COVID-19 and other diseases^8^, development of an all-human LF model using easily accessible cell sources also could provide a valuable preclinical tool for evaluation of vaccines and adjuvants.

In the present study, we therefore set out to explore whether human organ-on-a-chip (Organ Chip) microfluidic culture technology that has been previously used to recapitulate both normal physiology and disease states with high fidelity in multiple other human organs (e.g., lung, intestine, kidney, bone marrow, brain, etc.)^9–16^ can be leveraged to meet this challenge. Here, we show that primary B and T lymphocytes isolated non-invasively from the blood of multiple human donors spontaneously self-assemble into LFs that express AID and CXCL13 and support plasma cell differentiation when cultured within an extracellular matrix (ECM) gel contained in one channel of a two-channel Organ Chip. These studies revealed that dynamic fluid flow made possible by microfluidic culture both supports LF formation and prevents autoactivation previously reported in high-density cultures of human B cells^17^. Human LF Chips additionally containing autologous dendritic cells (DCs) display antigen-specific IgG production as well as secretion of clinically relevant cytokines when vaccinated with a commercially available, split virion trivalent influenza vaccine, and this same model is able to detect germinal center enhancing activity of a low cost squalene-in-water emulsion (SWE) adjuvant that could have great value for vaccination in low resource nations.

## RESULTS

### Self-assembly of T and B lymphocytes into LFs on-chip with minimal autoactivation

We used a microfluidic Organ Chip containing two channels separated by a porous membrane^18^ in which human T and B lymphocytes isolated from blood apheresis collars were cultured at a high density (1.5-2 × 10^8^ cells/ml; 1:1 ratio) within an ECM gel composed of Matrigel and type I collagen. The gel filled the lower channel of the device and was supplied with oxygen and nutrients through continuous perfusion of culture medium through the upper channel (Figure 1a; Supplementary Figure 1a). We chose this high culture density because high-density lymphocyte suspension cultures have been shown to be more predictive of human clinical responses than lower density cultures^19^. Human T and B lymphocytes commonly only form small cell aggregates when co-cultured in suspension or on membranes,^20^ and we observed similar behavior when we cultured unstimulated, patient-derived T and B cells at high density under static conditions in the same ECM gel (Figure 1b). In contrast, when these patient-derived T and B-lymphocytes were cultured under dynamic flow on-chip for 3-4 days, they spontaneously self-assembled into much larger 3D multicellular aggregates resembling small LFs (Figure 1b, Supplementary Figure 1b).

**Figure 1.**
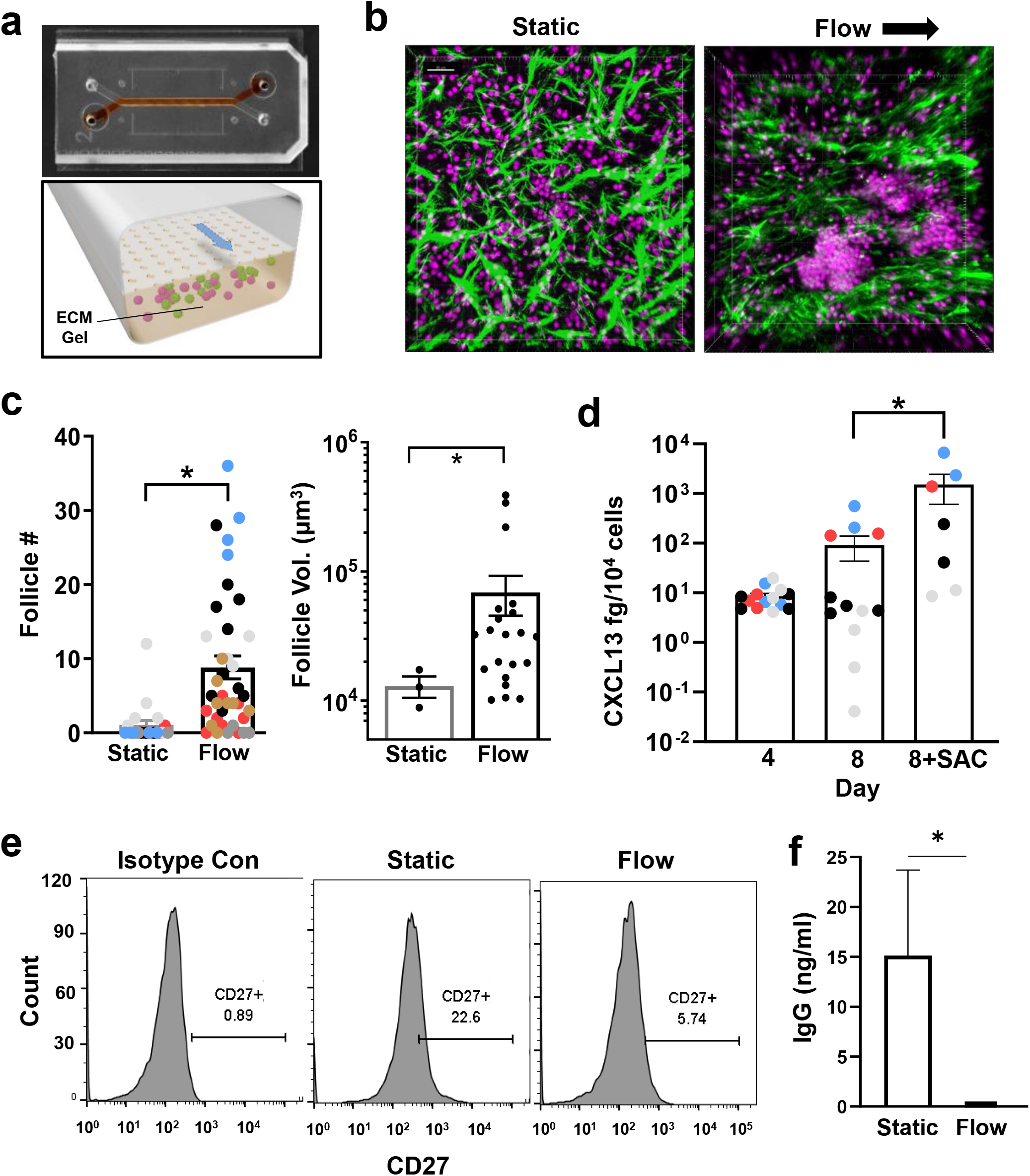
Perfusion induces follicle formation in the human LF chip. **a)** Photograph (top) of the 2-channel Organ Chip device used to create the human LF Chip with red dye filling the lower microchannel and a schematic of a cross section of the device (bottom) showing how the lower channel is filled with an ECM gel containing human lymphocytes, which are fed through the porous membrane separating the channels by medium that is flowed through the upper channel. **b**) Second harmonic images of ECM fibrils (green) combined with fluorescence images of Hoechst stained nuclei (magenta) of human lymphocytes growing at high density within ECM gels maintained within the lower channel of the LF Chip either under static conditions (Static) or with dynamically perfusion (Flow; arrow indicates direction) (bar, 30μm). **c)** Quantification of LF number (Follicle #) and volume (Follicle Vol.) in ECM gels that were cultured statically (Static) or with active perfusion (Flow) in Organ Chips for 4 days. Different colored data points represent follicle numbers (left) per field of view from >2chips Chips with cells from 6 different donors; *, *p* < 0.05. The volume for each follicle observed under Static or Flow is reported for one representative donor (right). Similar differences in follicle size were observed in two additional donors. **d)** Secreted CXCL13 protein detected within the effluent of the LF Chip cultured for 4 or 8 days in the presence or absence of SAC antigen using a Simoa assay. Different colored data points represent results from 1-4 chips created with cells from 4 different donors; *, *p* < 0.05. **e)** CD27 expression on B cells assessed by flow cytometry in isotype controls (left), static culture (middle) and the perfused LF chip (right). Representative results displayed 1 chip from one donor, and similar results were obtained with 2 chips using cells from two different donors; the percentage of cells in the positive peak is indicated below CD27+ in each graph. **f)** IgG levels determined by ELISA and normalized for culture volume in static culture (Static) versus perfused LF chips (Flow) in 2 chips each from 2 donors.

Second-harmonic microscopic imaging of these cultures revealed that dynamic perfusion of the chip resulted in realignment of matrix fibrils along the flow lines within the ECM gel whereas the fibrils formed larger aggregates and were oriented randomly in the static gels (Figure 1b). Interestingly, this force-induced structural reorganization of the ECM was accompanied by a significant increase in the number and size of the LFs compared to static ECM gels after 3 days of culture when analyzed in chips created with cells from multiple different human donors (Figure 1b,c; Supplementary Figure c,d). In the static gel cultures, LF formation was only observed in ∼25% (1 of 4) of the human donors, whereas 100% (6 out of 6) of the perfused microfluidic chip cultures formed LFs, and these were also significantly larger in volume (Figure 1c). However, the total number of LFs did not increase beyond 4 days and appeared to slightly decrease by 7 days of culture in the absence of antigenic activation (Supplementary Figure 1d). T and B cells also were found to be in close contact within the follicles, forming cell-cell contacts on-chip with polarized expression of the T cell co-receptor CD3 (Supplementary Figure 1e). Culture of lymphocytes on ECM-coated plates induces a similar tissue-like polarization in circulating T cells^19, 21^. Similarly, we found T cell polarization to be significantly increased in the LF Chips containing cells in 3D ECM gels whether maintained under static or perfused conditions compared to cells in conventional planar 2D culture lacking any ECM (Supplementary Figure 1e).

Lymph nodes are formed because of a positive feedback loop between lymphoid tissue inducer and organizer cells leading to production of CXCL13, CCL19, and CCL21 chemokines that promote recruitment of lymphocytes^22^. While the cellular and molecular cues that lead to ectopic LF formation are not fully defined, CXCL13 is key requirement for LF assembly^26^ and it is often used as a biomarker of ectopic LF formation^4^. CXCL13 levels in plasma also correlate with both LF formation and vaccine responses in humans^23, 24^. Moreover, CXCL13 and its receptor CXCR5 are key genes in the transcriptomic signature exhibited by tertiary lymphoid organs, which contain ectopic LFs^4^. Importantly, we found that the human lymphocytes produce CXCL13 under baseline conditions when cultured for 4 to 8 days on-chip, and its expression increased significantly by day 8 when the LF Chips were exposed to the bacterial *S. aureus* Cowan I (SAC) antigen (Figure 1d).

Autoactivation of B cells has been reported to be a challenge in high density cultures^17^, and so we expected that the increased numbers of cell-cell contacts we observed in our LF chips might lead to autoactivation as well. However, we did not observe significant autoactivation of B cells in our perfused LF chips, even though the same cells became autoactivated when cultured at the same high density under static conditions, as seen by an increased percentage of cells that express CD27 compared to cells cultured on-chip under flow or an isotype control (22.6% vs. 5.7% vs. 0.9%, respectively) (Figure 1e). To further assess autoactivation, naive B cells isolated from blood apheresis collars using CD27 depletion (Supplementary Figure 2a) were combined with T cells and either placed in static cultures or integrated into the microfluidic Organ Chips. We observed spontaneous class switching in static cultures as seen by production of high levels of IgG (Figure 1f), which was again consistent with the cells autoactivating. Importantly, however, we did not observe any class switching in the perfused LF chips under these baseline conditions (Figure 1f). Similarly, IgM levels were extremely high in static cultures, but not in the perfused LF chips (Supplementary Figure 2b).

### Ectopic LFs induce AID expression on-chip

B cells in the blood are a minor population of peripheral blood mononuclear cells (PBMCs) compared to T lymphocytes, and they are mostly naïve with about 70% appearing as IgD^+^ CD27^-^ cells when analyzed by FACS analysis, while tonsils that contain multiple LFs have a high number of B cells, but only about 40% of them are naive IgD^+^ CD27^-^ cells^25, 26^ (Figure 2a). Importantly, the unstimulated B cells within the LF Chips retained the naive phenotype of PBMCs, again with about 70% of cells appearing IgD^+^ CD27^-^ (Figure 2a).

**Figure 2.**
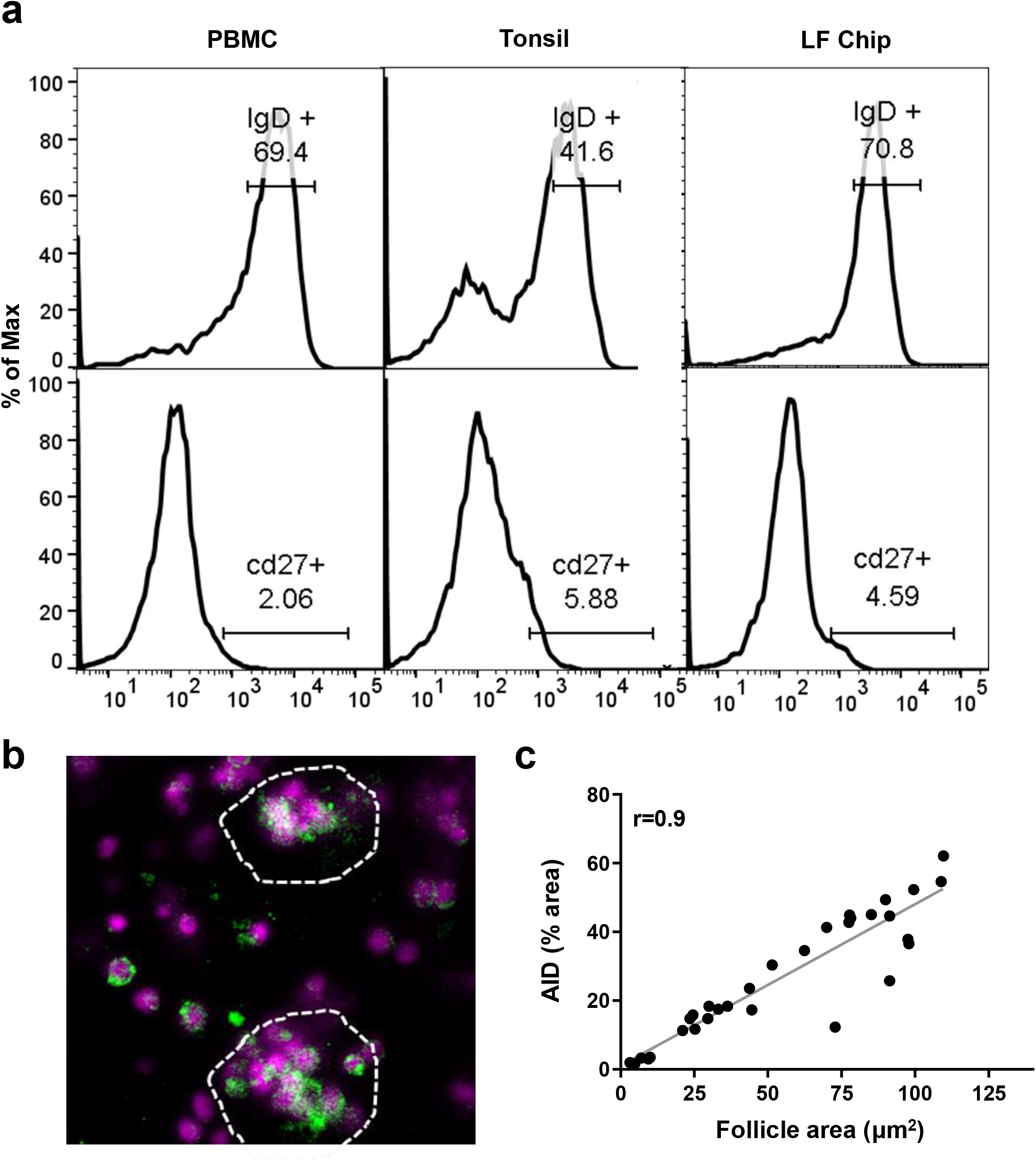
B cells remain quiescent yet express AID in LFs formed on-chip. **a)** Representative flow cytometric characterization of B cells stained for IgD and CD27 in the initial PBMC sample (PBMC; n= 3) compared with cells from explanted tonsils (Tonsil; *n* = 2), or cells cultured in the LF chip for 4 days (LF Chip; n=3). The percentage of cells in the positive peak are indicated above the gate drawn on the histogram. **b)** Representative confocal immunofluorescence micrograph showing AID expression (green) in B cells cultured in the LF chip for 4 days (similar results were obtained with 4 donors); Hoechst stained nuclei in the lymphocytes are shown in blue. **c)** Quantification of AID expression levels in individual follicles measured as % of the projected area of each LF expressing AID plotted as a function of LF size (Follicle area) in perfused Organ Chips cultured for 4 days. Each data point represents one follicles and follicles from 2 LF chips each created with cells from 4 donors were pooled for this analysis. Follicles were identified as regions of interest (ROI) by particle analysis in ImageJ. The percentage of the total ROI area that was positive for CD138 (y-axis) was measured and plotted against total follicle area (x-axis), Correlation analysis between was performed in Graphpad Prism (r=0.9).

Surprisingly, despite retaining a naïve phenotype, B cells within the ectopic LFs that self-assembled on-chip expressed the AID enzyme (Figure 2b), indicating the acquisition of follicular functions that are not normally present in circulating B cells. AID expression is important because it mediates the critical Ab class switching response within germinal centers in the lymph node *in vivo*.^5^ While we detected AID expression in B cells that primarily appeared within the LFs, some single cells within the organ chip were also AID^+^, as detected using confocal immunofluorescence microscopy (Figure 2b; Supplementary Figure 3a). However, when we quantified the number of AID^+^ B cells within LFs, we found that AID expression increased linearly with follicle size until the aggregates reached about 120 um^2^ in projected area (Figure 2c) but saturated thereafter (Supplementary Figure 3b). Lymphocytes cultured within the perfused Organ Chips also exhibited greater total AID expression as they promoted more LF formation compared to static ECM gel cultures (Supplementary Figure 3c).

. Flow cytometry was used to compare the number of B cells expressing IgM or IgD, follicular T helper cells (Tfh) as defined by co-expression of both CXCR5 and PD-1, and B cell expression of CXCR5 in the LF Chips compared to conventional static culture in a plastic dish (Supplementary Figure 4a). IgM^+^ cells in the LF Chips are composed mainly of naive B cells, whereas the IgG^+^ cells represent a small population of previously activated, class switched memory cells. This analysis revealed that compared to static 2D cultures containing B, T, and DC cells, there was a significant increase in the total cellularity of the unvaccinated LF Chip, including survival of naïve B cells, in 2 out of the 3 donors tested (Supplementary Figure 4b). Flow cytometric analyses also revealed that most of the B cells in the naïve LF chips express the CXCR5 receptor for CXCL13, and a fraction of the CD4 and CD8 T cells are also CXCR5^+^ (Supplementary Figure 4a). Thus, the CXCL13-CXCR5 signaling pathway appears to be activated under baseline conditions in B cells cultured in the LF Chip.

### Class switching and plasma cell differentiation

Naive B cells stimulated with IL4 and anti-CD40 Ab have been shown to undergo antibody class switching^27^, and thus, we used these stimuli to further characterize B cell functionality in the LF chips. When we perfused the LF chips with IL-4 and this CD40 ligand, IgG was detected in the chip outflows whereas it was not detected in unstimulated chips (Figure 3a), confirming that B cells are functional and capable of class switching in these Organ Chip cultures. Importantly, we also observed formation of CD138^+^ plasma cells and their organization within large clusters in many follicles on day 7 in the stimulated LF chips (Figure 3b). While CD138 expression could be detected in some single cells, again it was preferentially expressed by cells that clustered within the self-assembled LFs (Figure 3c), thus mimicking plasma cell development and differentiation that occur in LFs *in vivo*^28, 29^.

**Figure 3.**
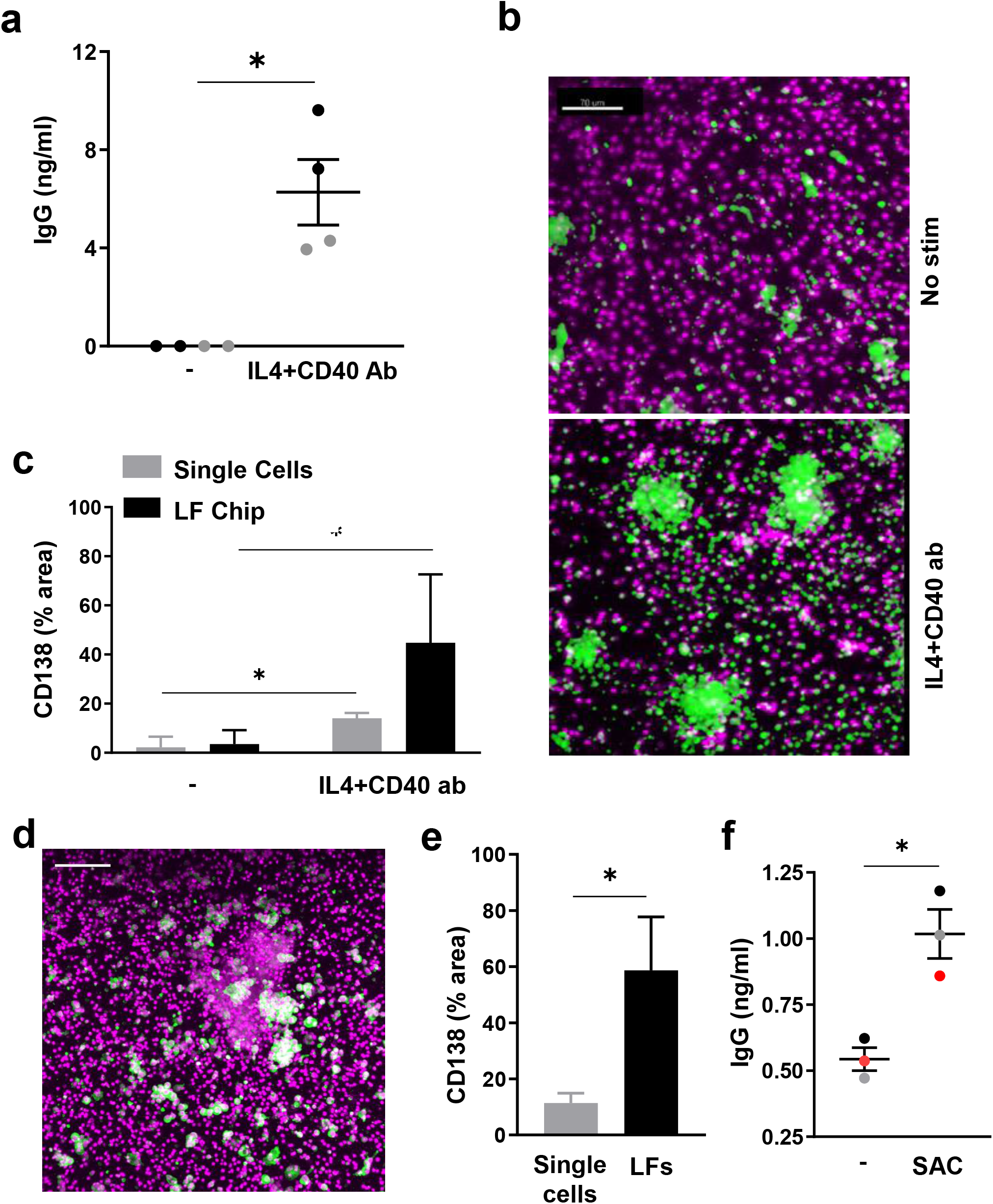
B cells exhibit class switching and undergo plasma cell formation in the LF chip. **a)** Total IgG production measured in the effluents of LF chips when engineered with naïve B cells and bulk T cells after 6 days culture in the presence or absence (-) of IL4 and anti-CD40 Ab Each dot indicates results from an individual chip (n=2) created with cells from two donors (black and gray); *, *p* < 0.05. **b)** Immunofluorescence micrographs showing cells in unstimulated LF chips (No stim.) or chips treated with IL4 and anti-CD40 Ab stained for CD138 (green) and nuclei (magenta); similar results were obtained with cells from 3 different donors. **c)** Quantification of CD138 expression in single cells (gray bars) versus cells located within LFs (black bars) in the same LF chips. Error bars indicate standard deviation based on analysis of 5 randomly selected fields from 1 donor, and similar results were obtained with LF Chips containing cells from 3 different donors; *, *p* < 0.05. **d)** Immunostaining for CD138 (green) and nuclei (magenta) in SAC-treated LF Chips (similar results obtained with 3 donors; bar 100μm). **e)** CD138 levels measured as a % of projected area labeled for CD138 in lone cells (Single Cells) versus cells in follicles (LFs) within ECM gels in perfused Organ Chips. Results shown are from 5 randomly selected fields from 1 LF Chip created with cells from one donor, and similar results were obtained with cells from 3 different donors; *, *p* < 0.05. **f)** Total IgG levels measured in the effluent of LF Chips 3 days after treatment with SAC. Each colored dot indicates results from one chip from each of 3 different donors; *, *p* < 0.05.

As ectopic LFs can be induced to form as a result viral^30^ or bacterial^31^ infections, we next explored if these ectopic LF Chips can be used as preclinical tools to study adaptive immune responses to pathogens by using the SAC antigen to mimic the presence of dead bacteria. Exposure to this bacterial antigen again induced CD138^+^ plasma cell formation to a much greater degree in the LFs that formed on-chip compared to nearby single cells (Figure 3d,e), and importantly, this was accompanied by robust production of IgG (Figure 3f). The LF chip therefore provides a way to model of ectopic LFs that self-assemble, form plasma cells, undergo class switching, and generate polyclonal IgG Abs in response to bacterial antigens.

### The LF Chip recapitulates human vaccine and adjuvant responses

Existing preclinical models used for testing of vaccines involve static 2D cultures or animal models that do not faithfully mimic human responses. Even more concerning is that the animal model that is closest to human – non-human primates – is currently in short supply and encumbered by ethical concerns^8^. Protective immunity induced by vaccination requires plasma cell production of antigen-specific antibodies, and vaccination-induced formation of ectopic LFs can contribute to this response^32^. Thus, we next explored with the human LF Chip can be used as study human vaccine responses.

As vaccine antigens are presented by DCs in the draining lymph node *in vivo*, we differentiated autologous human DCs from monocytes and included them in the ECM gel along with B and T cells as 2% of the total population. There have been reports suggesting that influenza vaccination can be performed *in vitro* using peripheral blood derived 2D cultures; however, later studies found that most of the responders were previously exposed to influenza^36^. To explore the utility of the human LF Chip as a preclinical human *in vitro* testing platform that can be used to derisk adjuvants as well as vaccines, we prepared 2D cultures and LF Chips from the same cellular pool containing an equal ratio of T and B cells with 2% DCs. Responses of the chips to vaccination with split virion influenza alone or combined with SWE adjuvant were analyzed using cells from 3 separate donors.

Unfortunately, we found that the sensitivity of commercially available plate-based ELISA methods and hemagglutination inhibition assays commonly used to detect anti-influenza hemagglutinin (HA) antibodies in human serum is too low for in vitro studies, even when using primary tonsil cultures vaccinated with the widely used split virion trivalent influenza vaccine Fluzone (Supplementary Figure 5a). To overcome this challenge, we modified a high sensitivity digital ELISA assay^33^ to detect influenza virus strain-specific antibodies and confirmed that it could detect high levels of anti-HA antibodies generated by lymphocytes in 2D tonsil cultures when stimulated with vaccine (Supplementary Figure 5b vs. 5a).

When we used this more sensitive assay, we found that human LF chips demonstrated improved antibody responses compared to 2D cultures lacking ECM and these responses were further enhanced by inclusion of the SWE adjuvant (Figure 4a). When tested in 2D cultures, only minimal anti-HA Ab production was observed, with only 1 donor and 2 out of 7 wells responding when vaccinated with inactivated H5N1 alone, and while addition of the SWE adjuvant resulted in responses in 2 out of 3 donors (and 4 of 7 wells), there was only a marginal improvement in the Ab response (Figure 4a). While vaccination of human LF Chips with inactivated H5N1 alone led to anti-HA Ab production in all 3 donors, the levels were quite variable and only 4 of 12 chips exhibited Ab production. In contrast, combined inoculation of the chips with both the vaccine and the SWE adjuvant stimulated high levels of anti-HA Ab production in all donors and in 7 of 11 LF chips (Figure 4a). This enhancement in antibody responses also was accompanied by an increase in follicle size and number in both donors tested suggesting that increased LF formation enables better Ab responses in vitro (Figure 4b,c). Moreover, a greater number of LFs were maintained in culture at 9 days after vaccination with SWE adjuvant compared to antigen alone (Supplementary Figure 6). While inactivated H5N1 lead to a modest increase in plasma cell number, SWE significantly enhanced the number of CD138+ plasma cells as quantified by the CD138+ area in each field of view (Figure 4d). Taken together, these studies show that the human LF Chip is more effective at both promoting LF formation and generating antigen-specific Ab responses than conventional 2D cultures, and it can be used to assess human vaccination responses as well as the modulating effects of potential adjuvants *in vitro*.

**Figure 4.**
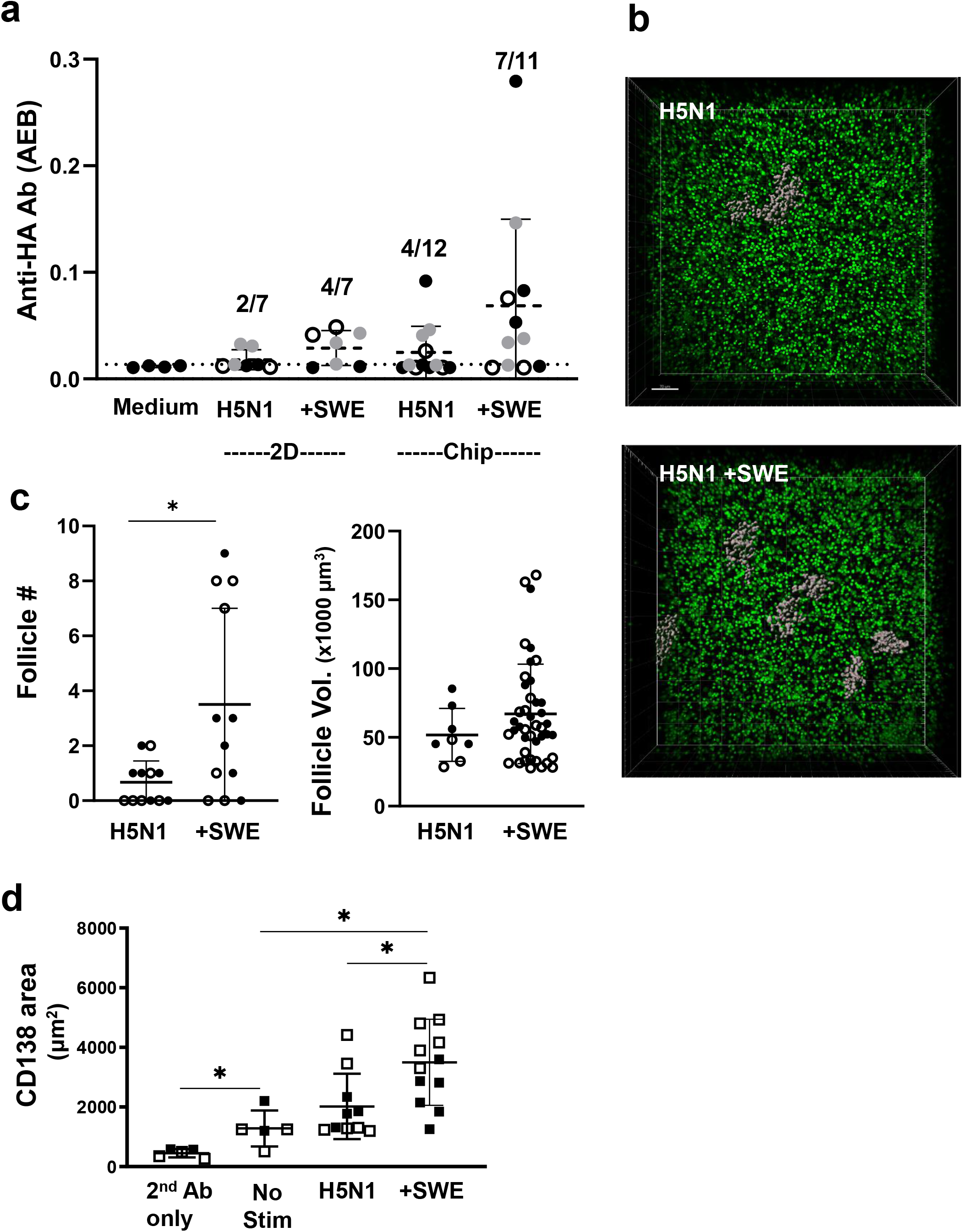
Vaccine and adjuvant induced antibody production and follicle formation in the human LF chip. **a)** Anti-HA Ab signal detected in the effluents of the LF Chips (Chip) or in the medium of cells maintained in static 2D culture (2D) for 9 days using the Simoa digital ELISA when unstimulated (Medium) or stimulated with H5N1 antigen alone (H5N1) or with SWE (+SWE). Each data point indicates one well or chip; different colored points indicate chips or wells from 3 independent donors. The fraction of total samples that exhibited anti-HA Ab production is indicated in the text above the bars. Simoa values are presented as average enzyme per bead (AEB). **b)** A 3D confocal microscopic stack view showing pseudocolored follicles (grey) and cell nuclei (green) present within ECM gels cultured for 3 days within a perfused LF Chip when vaccinated with a split virion H5N1 influenza vaccine in the absence (H5N1) or presence of SWE adjuvant (H5N1 + SWE); bar, 50 μm. **c)** Graph showing quantification of the number (left) and size (right) of follicles observed in LF Chips generated with cells from two different donors (open and filled circles); *, *p* < 0.05. Each data point indicates one field of view (follicle #) one individual follicle (follicle vol.). **d)** Graph showing quantification of the CD138+ area observed in LF Chips generated with cells from two different donors (open and filled squares); *, *p* < 0.05. Each data point indicates one field of view.

Finally, we explored whether we could generate clinically relevant vaccination response sin the LF Chips by inoculating them with Fluzone. Using this method, we found that LF Chips generated with cells from 8 different human donors all produced anti-HA antibodies at detectable levels, however, we detected the two patient subsets when we compared these results to Ab levels present in the supernatant of a vaccinated tonsil culture (Figure 5a), which produced a high anti-HA Ab signal in the digital ELISA (Supplementary Figure 5b). LF Chips from one patient group responded by producing similar levels of anti-HA IgG in the LF Chip as in the tonsil preparation whereas the other were low responders (Figure 5a). Abs were first detectable in the chip effluents about 5 days after immunization and their levels increased significantly over the following week in both the low and high responder groups (Figure 5b). Plasma cells also were detected at 5 days after immunization in the vaccinated LF Chips made with cells from donors that exhibited high levels of Ab production (Figure 5c). Importantly, this is the same time when circulating plasma cells can first be detected in the blood of vaccinated individuals^34^. Also, as expected based on past work showing the importance of antigen-presenting DCs for generation of a vaccination response^35^, there was a significant reduction in levels of anti-influenza HA-specific IgG (Figure 5d) and CXCL13 (Figure 5e) when DCs were not included in the LF chips. Interestingly, levels of CXCL13 measured on-chip 5 days post vaccination were also highly predictive of anti-influenza Ab responses in the high responder group (Figure 5f).

**Figure 5.**
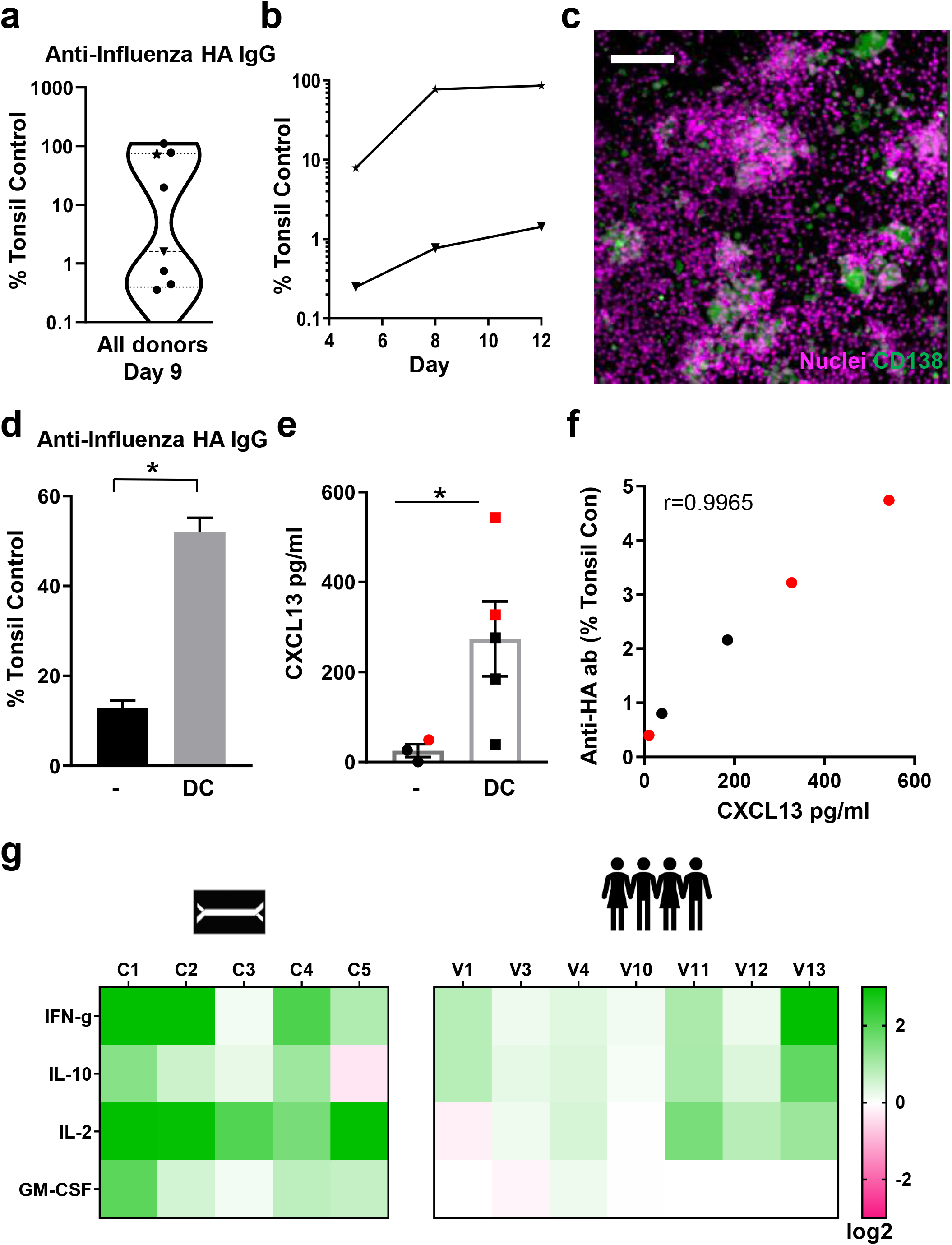
Influenza vaccination *in vitro* in the human LF chip. **a)** Violin plot of anti-HA IgG levels that are specific to the Brisbane 59 H1N1 strain (Anti-HA IgG) in the effluent of LF Chips 9 days after vaccination with Fluzone presented relative to levels measured in a vaccinated culture of tonsillar cells (Tonsillar Control), as detected using a digital ELISA. Each data point represents average of 2-6 chips created with cells from one donor (total 8 donors tested). **b)** Time course of anti-HA IgG secretion measured in chip effluents over 5 to 12 days of culture using LF Chips containing cells from representative donors from the high (★) and low (▼) Ab producer groups shown in **a**. **c)** Immunofluorescence micrograph showing CD138 staining (green) in a Fluzone-stimulated LF Chip containing cells (nuclei, magenta) from a high anti-HA Ab producer (similar results were obtained with 3 different high Ab producer donors; bar, 100μm). **d)** Anti-HA Ab levels that are specific to the Brisbane 59 H1N1 strain in the effluent of LF Chips with or without DCs, 9 days after vaccination, as detected by a digital ELISA, presented relative to levels measured in a culture of tonsillar cells. Mean levels from 3 replicate measurements from one chip generated from one donor are shown, and similar results were obtained in LF Chips created with cells from two different donors; *, *p* < 0.05. **e)** CXCL13 levels in the effluent of the LF chip with or without DCs, 5 days after vaccination, as detected by a digital ELISA. Each colored dot represents one chip with cells lined from two different donors (black and red dots); *, *p* < 0.05. **f)** Levels of anti-HA Ab specific to the Brisbane 59 H1N1 strain in the effluent of LF chips plotted relative to CXCL13 levels 5 days after vaccination. Results from 2 donors are shown with each colored dot representing an individual chip; an analysis of the correlation between CXCL13 and IgG levels are in Graphpad Prism is shown; r=0.99 **g)** Heat map of the fold change (Log_2_) in the levels of cytokines (IFN-γ, IL-10, IL-2, GM-CSF) measured using a digital ELISA in effluents of LF chips generated with cells from 5 different donors, 3 days after vaccination (C1-5) compared to unvaccinated chips, and to levels measured in peripheral blood from 7 individuals (V1, 3, 4, 10-13) 1 day after vaccination as compared to their prevaccination levels.

Finally, we quantified levels of four cytokines within the effluent of the vaccinated LF chips that are important for T cell expansion, survival, and helper functions (IFN-γ, IL10, IL2, GM-CSF), and compared their levels to those measured in the serum previously obtained in a study of human volunteers vaccinated with influenza vaccines. Serum was collected from volunteers before and one day after vaccination and frozen for further analyses. These studies showed that the responses of all four of these cytokines induced in the human LF chip by vaccinating with Fluzone are very similar to those observed in immunized human volunteers *in vivo* (Figure 5g).

## DISCUSSION

Animal models are the gold standard for advancing vaccine candidates and immunotherapeutics to the clinic. However, because the immune systems of animals are significantly different from humans, unpredicted toxicities or poor efficacies have resulted when vaccines and immunotherapies entered clinical trials^1^. For example, the severe cytokine syndrome that developed in patients enrolled in the Phase I clinical trial of the CD28 superagonist TGN1412 had not been detected in preclinical animal models^2^. High density cultures of PBMCs were later shown to predict high levels of cytokine release by TGN1412; however, it is not possible to use these static 2D cultures to study the how immune responses induce reorganization of lymphocytes in 3D that leads to LF formation and expansion, as well as associated Ab class switching, plasma cell formation, germinal center organization, and secretion of high affinity Abs. Humanized mice have been explored as a possible alternative to study vaccines *in vivo*, but they are severely immunocompromised, require use of human fetal tissues which has ethical implications, and they lack lymph nodes, which causes defects in class switching, affinity maturation and long-lived plasma cell formation^39^. Most concerning is that during the current COVID19 pandemic, non-human primates that have been used to test vaccine have become a scarce resource^8^, which could limit development of newer vaccines that might be required as new variants emerge.

In this study, we described microfluidic Organ Chips containing primary human blood-derived B and T cells isolated non-invasively from peripheral blood that spontaneously self-assemble into LFs that undergo Ab class switching, form plasma cell clusters, organize germinal centers, and secrete high affinity antigen-specific IgG when stimulated with antigen. Importantly, when we also integrated autologous DCs into these LF Chips, we found that they can be used to evaluate the efficacy of vaccines as well as adjuvants. This was demonstrated by vaccinating the chips in vitro using a commercial influenza vaccine and a low cost SWE adjuvant that is being developed for use in resource poor nations. It is important to note that others have studied cultured human tonsillar lymphocytes obtained from patients^40, 41^ or created putative 3D models of the lymph node^42, 43^ to conduct preclinical testing of vaccines. However, these experimental systems do not demonstrate lymph node-like biomarkers, *de novo* LF formation, or survival of plasma cells, and hence, cannot be used to study LF formation, physiology, or the underlying basis of human adaptive immune responses *in vitro* as the LF Chip can.

The formation of germinal centers is a hallmark of protective immunity against infectious agents and formation of germinal center-like structures in ectopic locations can predict vaccine^30, 32^ and immunotherapy^4, 44^ efficacy. However, ectopic LF formation also can result from pathological activation of the immune system, as observed in patients with autoimmunity ^45, 46^. There is currently no model of human LF formation where this process can be studied under controlled conditions in vitro. There was a recent report describing self-aggregation of tonsillocytes in static cultures^47^; however, tonsillar (and splenic) tissue is difficult for most labs to access as surgical intervention is required and visualization of higher order multicellular organization is difficult using this approach. In contrast, the in vitro Organ Chip approach we describe here results in self-assembly of LFs from peripheral blood-derived circulating immune cells, which are much easier to obtain. Thus, this is a major advantage of the LF Chip that is of particular value when trying to scale up this approach for preclinical testing of vaccines, for example.

Importantly, because we are able to control all components of the model and were able to observe LF formation over time *in vitro* without the addition of any chemical inducer or exogenous stromal or myeloid cells, we also were able to gain new insights into how follicle assembly is controlled. First, we were able to clearly demonstrate that the presence of B and T lymphocytes is sufficient to induce LF formation, which is consistent with recent studies of tertiary lymphoid organ formation^48, 49^. In addition, we discovered that dynamic fluid flow promotes LF formation, which appears to be mediated at least in part by flow-induced reorganization of ECM fibrils that could promote increased cell-cell interactions. Surprisingly, fluid flow was also required to avoid B cell autoactivation that occurs when these cells are cultured at a similar high density in static 2D cultures. These findings raise the intriguing possibility that dynamic perfusion of lymph nodes or tertiary lymphoid organs also may actively promote LF formation and growth, and perhaps even contribute to maintenance of a naïve state of these cells *in vivo*. This hypothesis is consistent with the observation that lymphatic vessel formation accompanies tertiary lymphoid organ formation and that lymph node formation can be inhibited by suppressing lymphatic vessel development^50, 51^. Physical forces and ECM reorganization underlie many developmental processes^52^ and the microfluidic LF Chip can be used to explore this mechanochemical mechanism in detail in the future.

Interestingly, we also discovered that high-density co-cultures of B and T cells in ECM gels cultured under fluid flow on-chip induce AID expression in B cells, and that the number of AID-expressing cells increases with follicle size. AID expression by these cells suggests that they might be in a transitory stage between circulating naïve B cells and activated germinal center cells, which have been recently described as pro-germinal center cells that are IgD^+^ AID^+^ CD27^-^.^53^ We also confirmed that there is a link between CXCL13 induction and follicle formation, which has been previously observed during vaccine responses in humans^23, 24^. In addition, CXCL13 and CXCR5 are highly represented in the genetic signatures of tertiary lymphoid organs as they are in our LF chips, again showing that this model faithfully recapitulates key features of ectopic LF formation *in vitro*^4^. In the current study, we focused on the development of a human in vitro model of ectopic LF formation using peripheral blood derived leukocytes and its potential use as an in vitro tool for assessing vaccine and adjuvant candidates in a more human relevant way. However, using this new experimental *in vitro* platform, it should be possible to define other key parameters that govern LF assembly, and also incorporate other relevant cell types (e.g., endothelial cells, stromal cells) to explore how they contribute the formation and function of LFs in the future.

Using the LF Chip, we found that challenge with 3 different types of immunological stimuli - IL4 combined with anti-CD40 Ab, SAC, or the clinically relevant Fluzone vaccine – all promote spontaneous LF development, plasma cell formation, and secretion of antigen-specific high affinity IgG. In contrast, current methods for plasma cell culture rely on extensive cytokine stimulation of memory B cells, stromal cell co-culture, and multiple rounds of manual handling^40, 54–56^. Treatment of human LF chips with IL-4 and anti-CD40 Ab led to robust expansion of plasma cells that were preferentially localized in LFs, and this did not occur in 2D cultures. It is important to note that other *in vitro* 3D models of the human lymph node have been explored in the past, such as the MIMIC system that mixes microbeads bound to T or B cells^42^, and the HuALN lymph node system that uses a continuously perfused mesh as a tissue-like support^43^. However, neither of these models produce detectable LF formation or robust plasma cell formation. In recently published tonsil organoids^47^, plasmablast formation can be observed in response to vaccination, but its formation and the physical and biochemical cues that control it cannot be studied in situ.

Testing new vaccines for emergent influenza virus strains is a huge burden on the healthcare system, as the vaccines must be repeatedly optimized because recombination between influenza virus strains is very common and influenza surface antigens can rapidly mutate. Influenza vaccines, which are currently, tested in ferrets, commonly only result in 30-60% protection in humans^57^. Accurate *in vitro* assessment of vaccines for new strains of influenza using primary human cells that generate plasma cells and undergo Ab class switching would therefore represent a major advance, in addition to significantly reducing the need to rely on animal models. In our studies, we were able to demonstrate successful vaccination in human LF Chips when inoculated with split virion H5N1 influenza particles, as measured by increased LF formation, plasma cell differentiation and anti-HA IgG production, and that we can detect augmentation of this response by addition of a low cost adjuvant (SWE) in vitro. In contrast, influenza vaccination alone or with SWE only induced modest anti-HA antibody responses in 2D cultures containing the same cell types, which do not support 3D structures. These data clearly show that the formation of 3D LFs within the microfluidic Organ Chip is a significant and novel advance over existing culture and animal models as it recapitulates vaccine-induced LF formation and production of specific high affinity IgG Ab in human-derived immune cells *in vitro*. Importantly, when we tested the utility of the human LF chip by vaccinating chips made from 8 independent donors with Fluzone, we detected donor-specific responses with formation of plasma cells in the high-responding donors who also produce cytokine biomarkers similar to those seen in vivo.

Thus, the LF Chip may provide a valuable preclinical tool that can overcome the limitations of humanized mice and even non-human primates to inform clinical vaccine trials design. In support of this possibility, IFN-γ was upregulated in the LF chip on vaccination as it is in humans, and this has been shown to be a key biomarker of vaccine efficacy^58^. As we used bulk B and T cells in the human LF Chips that were vaccinated, it is possible that the high responder cells had a preexisting memory for the Fluzone antigens. However, we chose this strategy as it replicates the administration of seasonal influenza vaccines in human patients who have had varying degrees of past exposure to the vaccine and closely related cross-reactive viral strains. Finally, the high sensitivity digital anti-HA ELISA described here can complement traditional methods of quantifying anti-influenza antibodies, which can predict protection from influenza in seropositive donors, but are severely limited in sensitivity.

In summary, this is the first reported *in vitro* model that supports formation of human LFs with functional germinal centers similar those found in secondary and tertiary lymphoid organs *in vivo*. In addition, we demonstrated that human LF Chips containing B and T cells along with autologous antigen-presenting DCs can be used to test human immunization responses to vaccines and adjuvants in vitro. In contrast to other culture models, the human LF Chip also can be used to study *in vivo*-like ectopic LF formation as well as antigen-induced immunological responses, and all of this is done using primary human blood cells collected non-invasively. Key improvements as compared to humanized mice and other existing *in vitro* systems used to study adaptive immune responses include 3D follicle formation, expression of AID and CXCL13, plasma cell formation, antigen-specific high affinity IgG Ab generation, and production of clinically relevant cytokines in response to multiple antigens, including a commercial influenza vaccine. The human LF Chip enables the longitudinal study of all these responses within a single device during the course of an experiment, and many of the processes can be observed over time *in situ* using high-resolution imaging. These microdevices are also patient-specific and thus, they can recapitulate donor variability in antibody responses to vaccination as seen in our studies with Fluzone. Taken together, these findings suggest that the LF Chip potentially may be useful as a more human relevant preclinical tool for assessing the efficacy and safety of vaccines, adjuvants, and immunotherapeutic drugs in a patient-specific manner.

## METHODS AND MATERIALS

### LF Chip Culture

Apheresis collars, a by-product of platelet isolation, were obtained from Brigham and Women’s Hospital under the approval obtained from the Institution Review Board at Harvard University. PBMCs were isolated by density centrifugation using Lymphoprep (StemCell Technologies, 07801), and magnetic beads were used for negative selection of bulk B cells (StemCell Technologies, 19054) or naïve B cells (17254), T cells (17951) and monocytes (19058). Isolated cells were counted and directly seeded on chip or frozen in Recovery Cell Culture Freezing Medium (Gibco, 12648-010). Monocytes were differentiated into DCs as previously described^59^. Briefly, 1 × 10^6^ monocytes were cultured in complete RPMI (RPMI 1640, Gibco, 72400-047), 10% Fetal bovine Serum [FBS, Gibco, 10082-147], 1% Penicillin/Streptomycin (Gibco, 15140122) and 400 ng/ml GM-CSF (Mitenyi Biotec, 130-095-372) and 250ng/ml IL-4 (130-094-117) for 5-6 days with 50% of the medium being replaced with fresh medium and cytokines every 2-3 days.

Organ Chips fabricated as previously described^18^ or obtained from a commercial vendor (Emulate Inc., Boston, MA) that contain two parallel channels separated by a porous membrane (7 *μ*m pores) were first perfused and incubated with 1% 3-Aminopropyltrimethoxysilane (Sigma, 281778) in ethanol for 1 hour and then incubated in an 80°C oven overnight. Cells were then suspended (1.5-2 × 10^8^ cells/ml) in complete RPMI containing Matrigel (60%; Corning, 356234) and type I collagen (15%; Corning, 354249) maintained on ice, and the viscous solution was then introduced in the bottom channel of the 2-channel Organ Chip. After the ECM was allowed to gel for 30 minutes in a 37°C cell culture incubator, the top channel of the chip was filled with medium and channel entry and exit ports were plugged with 200 *μ*l tips containing medium. The following day chips were perfused with RPMI medium containing at 60 *μ*l/hr using peristaltic pumps (Cole Parmer) or automated Zoe Organ Ohip instruments (Emulate Inc.) in a 37°C cell culture incubator following the manufacturer’s instructions. Both chip sources and perfusion systems produced similar results.

### 2D cell culture

To study immune response in conventional 2D culture and to compare the results to the Human LF chip, the lymphocyte mix was resuspend in medium at 2.25-3 × 10^6^ cells/ml and plated at 1ml/well in tissue culture treated 24 well plates or 0.5ml/well in 48 well plates. Vaccination was performed by adding vaccine with or without adjuvant in an equal volume for a total volume of 2ml/well in a 24 well plate or 1ml/well in a 48 well plate. To compare results between 2D culture and chip, cytokine and antibody results were normalized for volume and cell number.

### Activation of human LF Chip

To activate immune responses, chips were treated with immune activators by perfusing medium containing SAC (Sigma, 1:50000), IL4 (Miltenyi Biotec, 130-094-117, 400 *μ/*ml) and anti-CD40 antibody (BioXcell, BE0189, 1 *μ*g/ml]), or Fluzone (BEI resources, NR-19879, 10 *μ*l/ml) through the top channel of the device. Treatment with IL4 and anti-CD40 was initiated 2 days after seeding and SAC treatment was initiated 4 days after seeding (3 days of perfusion), and fresh medium was added to the inlet perfusion reservoirs as needed. In the Fluzone experiments, medium was recirculated for the first 5 days of treatment (i.e. effluents were added back to the inlet perfusion reservoir), and at day 5 and every 3 days thereafter, a 1:1 mix of effluent and fresh medium was used for perfusion. This was done to ensure that the cytokine milieu was maintained and to limit and taper the amount of Fluzone consumed by the experiment.

The following reagents were obtained through BEI Resources, NIAID, NIH: Fluzone® Influenza Virus Vaccine, 2009-2010 Formula, NR-19879 and HA1 Hemagglutinin (HA) Protein with N-Terminal Histidine Tag from Influenza Virus, A/Brisbane/59/2007 (H1N1), Recombinant from Baculovirus, NR-13411. Split virion H5N1was provided by Vaccine Formulation Institute (VFI, Belgium) ^60^. SWE and H5N1 were resuspended to an isotonic suspension using buffers provided by VFI. LF chips were vaccinated with 0.1 *μ*g/ml HA units and 10 *μ*l/ml SWE.

### Immunofluorescence Microscopy

To label B and T-cells for microscopy before seeding chips, lymphocytes were labeled with CellTracker dyes (Invitrogen, 1-2 μM). LF chips were fixed by filling the perfusion channel with 4% paraformaldehyde in phosphate-buffered saline (PBS, 1hr at room temperature, RT) and plugging the channel entry and exit ports with 200 μl tips containing fixative. To carry out immunostaining, chips were similarly incubated with primary antibodies diluted in PBS containing 1% FBS and Fc Block (1:20 dilution; Miltenyi Biotec, 130-059-901) overnight at RT, washed several times with PBS, and then incubated with secondary antibodies diluted in PBS containing 1% FBS overnight at RT, followed by one-hour incubation with Hoechst (Life Technologies, H3570, 1:1000) and several PBS washes at RT. ECM fibers were imaged using second harmonic imaging after fixation which allows the label-free imaging of biopolymers using a Leica SP5 confocal microscope. Follicle size and number were quantified using the surfaces function of Imaris (Bitplane) software (Supplementary Figure 1d). CD138 was quantified by analyzing particles in ImageJ.

### Quantification of T cell polarization

Samples were stained with a fluorophore labeled antibodies directed against CD3 (Biolegend, clone UCHT1), AID (Invitrogen, PA5-20012), or CD138 (Invitrogen, PA5-47395). Using ImageJ software, random cells were selected, and the center of mass of each cell was defined. Next, the center of mass of the CD3-positive fluorescent region was determined for each cell, and then the absolute distance between the center of the cell and the center of the CD3-stained region was used to define T cell polarization, with greater distances indicating more polarization.

### Flow cytometry

Cells were harvested from chips by blocking one port of the basal channel with a tip and manually pipetting Cell Recovery Medium (Corning, 354253 200 *μ*l/chip) through the other port to push the ECM and cells out of the channel and harvest them. The harvested ECM was incubated in the Cell Recovery Medium for one hour at 4°C. The released single cells were centrifuged at 300g for 5 min and resuspended in PBS for staining. Cells were first labeled with Live/Dead fixable dyes (1:1000 dilution; Invitrogen, L34963), followed by a 15 min incubation with fluorophore labeled antibodies and Fc Block (1:100 dilution) at 4°C. Cells were washed twice and then fixed with Cytofix (BD Biosciences, 554655) for 15 minutes at RT. Cells were centrifuged and re-suspended in PBS and stored at 4°C prior to cytometric analysis using a using LSRFortessa flow analyzer (BD Biosciences). Results were analyzed using FlowJo V10 software (Flowjo, LLC) using a volume of 100ul containing between 0.5 to 1 × 10^6^ cells. Antibodies to CD19 (#340437), CD27 (#655429), CD3 (#641406), CD4 (#555346), and CD8 (#347314) were obtained from BD Biosciences; antibodies to IgD (#348210) and CXCR5 (#356919) were acquired from BioLegend.

### Immunoglobulin and Cytokine Quantification

Total immunoglobulin levels were measured using ELISA [Bethyl Biolabs, E80-104 or Mesoscale discovery, K15203D). Influenza HA-specific IgG was detected using a modified version of a previously described digital ELISA assay^33^. Briefly, HA from Influenza A/59/Brisbane (BEI Resources, NR-13411) was conjugated to carboxyl-modified paramagnetic beads. Tonsils cultures (2.25-4 × 10^6^ cells/well of a 24 well plate) were immunized with 10ul/ml of Fluzone^61^ were used as positive controls. For detection of anti-HA antibodies, tonsil culture supernatants or chip effluents were incubated with the beads, followed by a biotinylated anti-human Ab [Life Technologies, A18821] and streptavidin-β-galactosidase and analyzed using Simoa HD-1 Analyzer (Quanterix). Tonsils were obtained from Massachusetts General Hospital (MGH) under an Institutional Review Board approved protocol. The digital ELISA was compared to a commercially available ELISA (Abcam, 108745) and found to have much higher sensitivity (Supplementary Figure 5). The sandwich digital ELISA was also used with antibodies from Biolegend directed against to IFN-*γ* (#507502), IL-7 (#501302), or IL-10 (#50680) and from R&D Systems against IL-2 (#MAB602), GM-CSF (#MAB615), or CXCL13 (#DY801). Secondary antibodies used in these studies against IL 10 (#501501) and IL-7 (#506601) were from Biolegend, and against IFN-*γ* (#MAB285), IL-2 (#MAB202), GM-CSF (#BAF215), CXCL13 (#DY801) were from R&D Systems. Protein standards for recombinant IFN-*γ* (#285-IF-100), IL-2 (#202-IL-010), IL-7 (#207-IL-005), IL-10 (#217-IL-005), GM-CSF (#215-GM-010), CXCL13 (#DY801) were all from R&D Systems.

Briefly, immune complexes were formed on beads in a 3-step format by 1) capturing target with antibody-conjugated beads, 2) binding with biotinylated detection antibody, and 3) labeling with streptavidin-β-galactosidase. In depth descriptions of Simoa assay development and performance have been previously reported^33^. Participant recruitment and sample collection followed an Institutional Review Board (IRB, Tufts University)-approved protocol (Study number 1410019). Informed consents were obtained from all participants in this study.

## Statistical Analyses

Student’s t-tests and tests for linear correlations were performed using Prism (GraphPad Software). Where distribution of data was not normal, the Mann-Whitney test was used. In all figures, * indicates *p* < 0.05 in all figures. The number of donors, chips/donor, and replicate experiments are described in the figure legends.

## ACKNOWLEDGMENTS

This research was sponsored by funding from the Defense Advanced Research Projects Agency under Cooperative Agreement Number W911NF-12-2-0036, the National Institutes of Health grant UG3HL141797, Bill and the Wyss Institute for Biologically Inspired Engineering (all to D.E.I.). The authors would like to thank D. Chou for helpful discussions, P. Sadow for providing surgically removed tonsils, and A. Sutherland for technical assistance.

## AUTHOR CONTRIBUTIONS

G.G. designed *in vitro* experiments with the help of G.M. G.G, G.M and P.P performed *in vitro* experiments and analyzed the data, working with D.E.I, who also supervised all work. Y.Z, M.S.K, A.M, D.C, Y.Z and J.L. assisted in performing experiments and analyzing the data. B.B, L.X, T.G, R.L and L.M. designed and conducted all high sensitivity digital ELISAs under the guidance of D.R.W. G.G, G.M, Y.Z and D.E.I. wrote the manuscript.

## SUPPLEMENTARY FIGURE LEGENDS

**Supplementary Figure 1.**
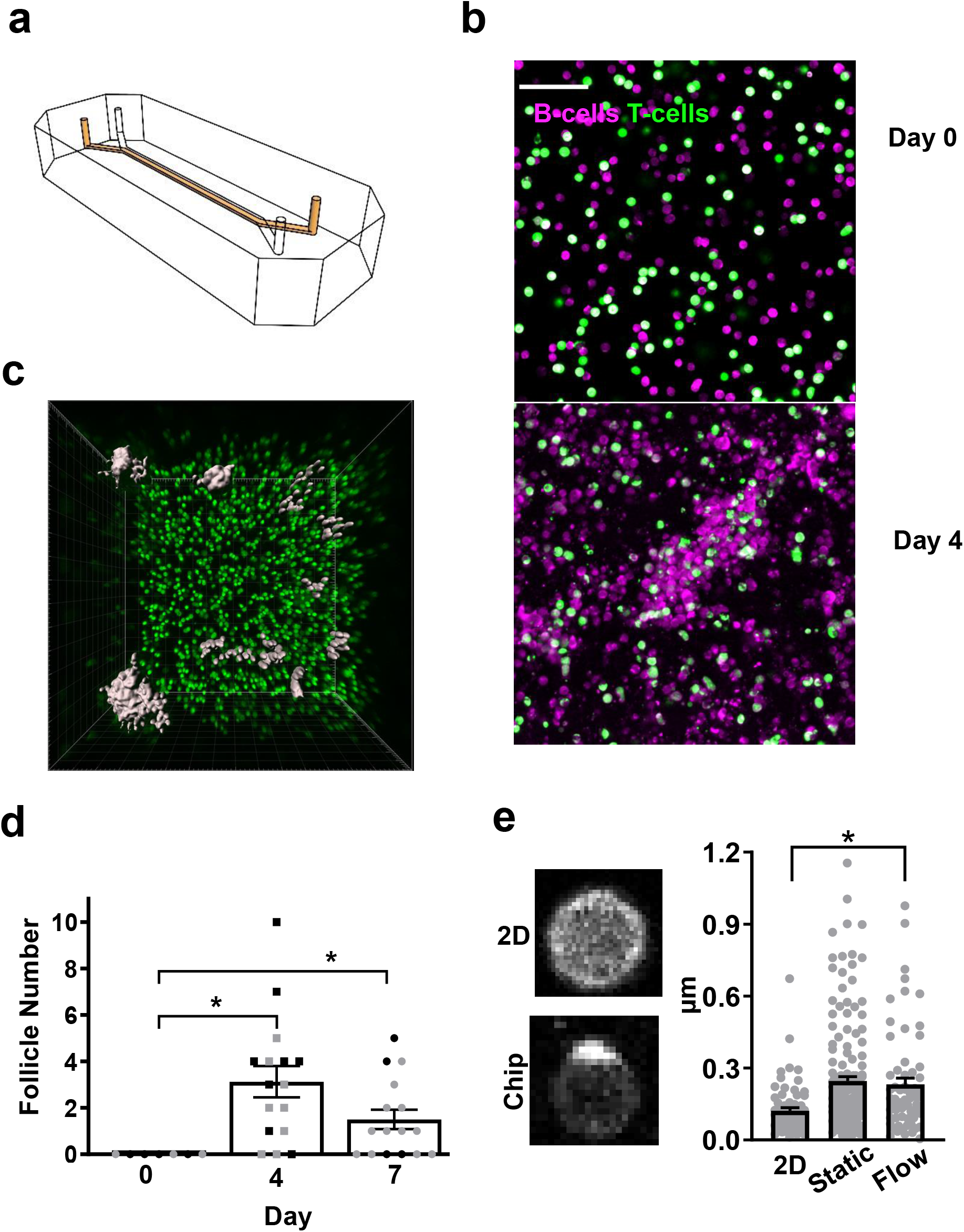
**a)** 3D drawing of Organ Chip with the channel containing the 3D ECM gel colored in orange; the parallel flow channel is not colored. **b)** Immunofluorescence micrographs showing CellTracker labeled human B (magenta) and T lymphocytes (green) cultured within an ECM gel in the LF Chip at day 0 and 4 (3 days after perfusion was initiated); bar, 50 μm). **c)** Representative surfaces seen on the LN chip after 4 days of culture visualized as surfaces with Imaris software. 1 field from a representative donor shown (similar results were obtained in 6 donors). **d)** Change in follicle number over time in 2 donors. Each dot represents one field. Donors are distinguished by color of the dot; *, p < 0.05. **e)** Confocal immunofluorescence micrographs of T cells in 2D culture (2D) versus perfused LF chip (chip) stained for CD3 (left), and quantification of T cell polarization carried out in 2D culture, static organ chips or the perfused LF chip (right) as described in Methods. Each dot is a cell and >64 cells were analyzed in all conditions. Results from one donor are shown, and similar results were obtained in LF Chips created with cells from three different donors; *, *p* < 0.05.

**Supplementary Figure 2.**
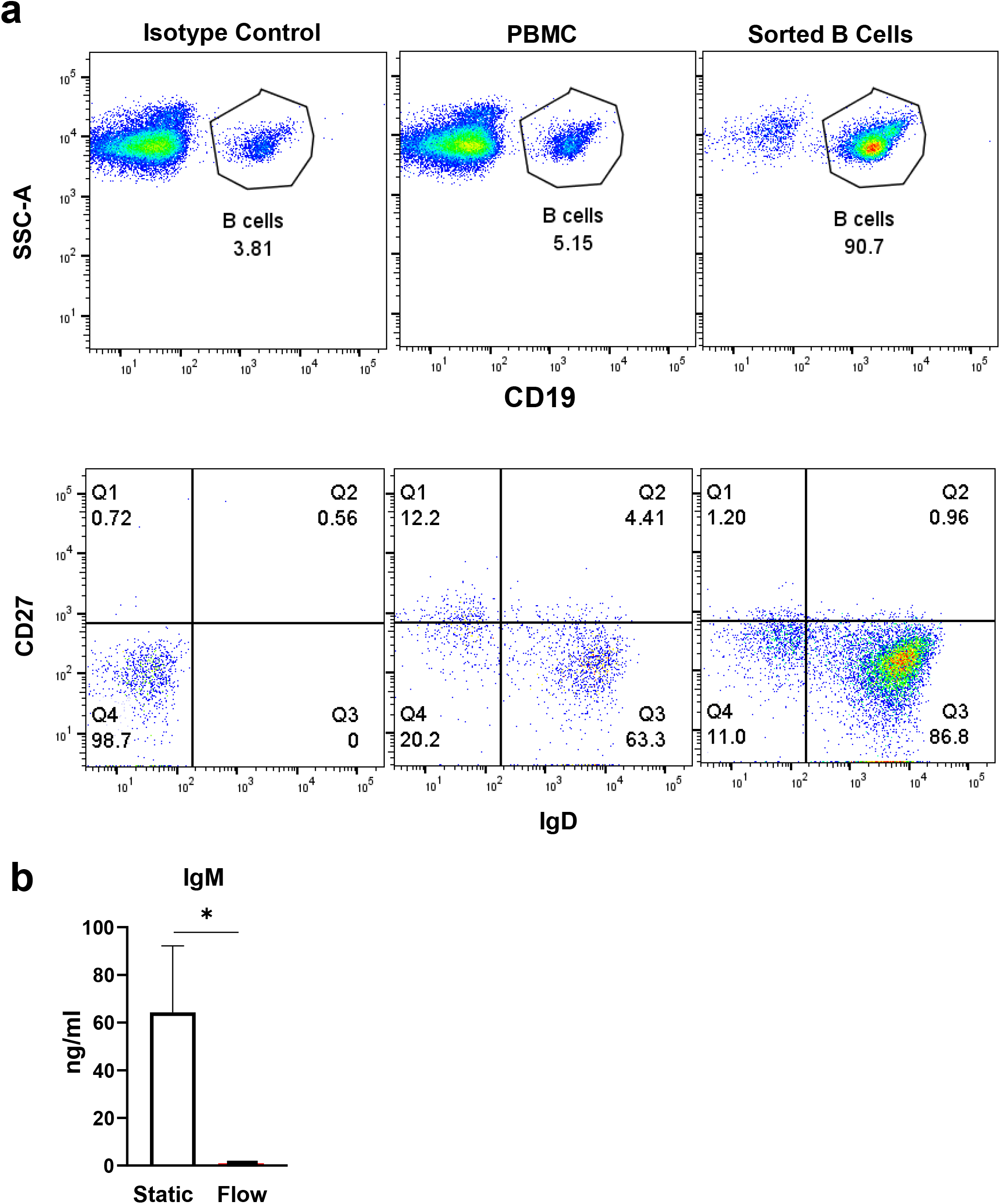
**a)** Purity of naïve B cells sorted from apheresis collars. 1 representative donor shown out of >6 donors tested. **b)** IgM secretion by unstimulated static cultures (Static) and perfused chips (Flow) after 7 days of culture. Mean IgM levels from 2 chips/condition from 2 donors (n=4) shown here. Error bars show standard deviation; *, p < 0.05.

**Supplementary Figure 3.**
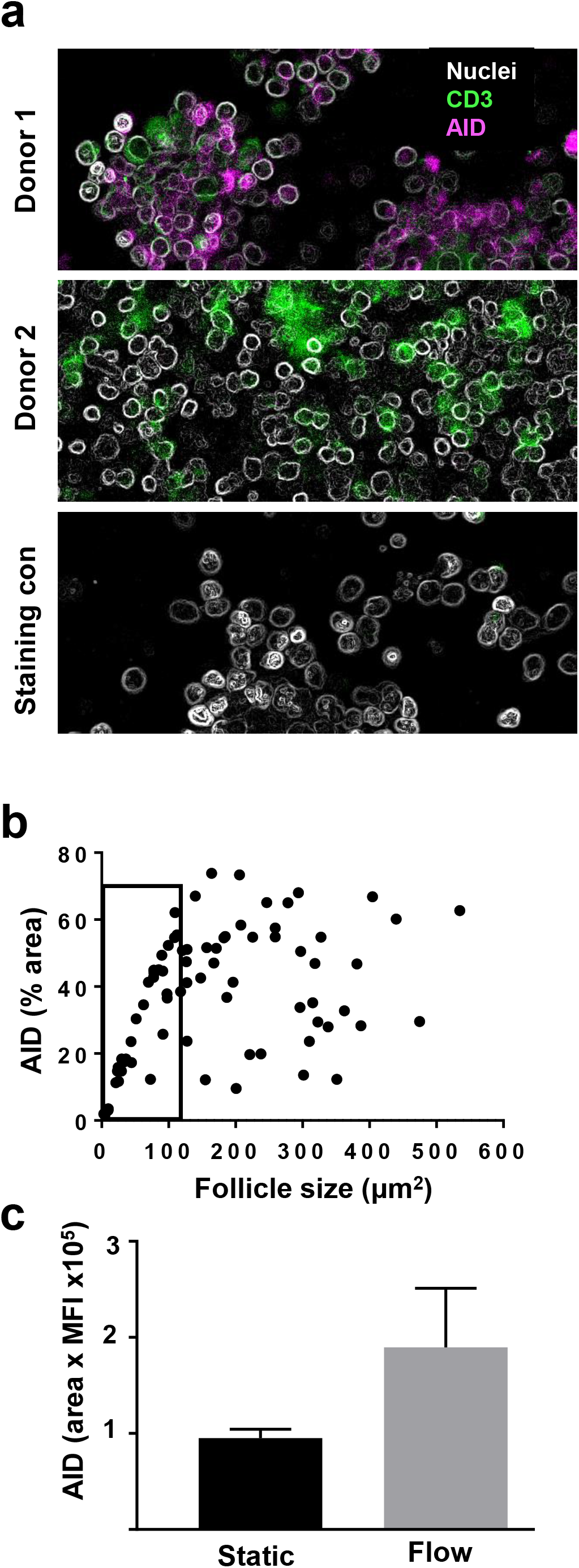
Optimization of AID staining for confocal microscopy. **A)** AID staining was tested on two donors after 4 days of static culture. **B)** AID expression in the LN chip. Each dot represents a follicle. Data pooled from 4 donors perfused on chip for 4 days. **C)** AID expression in a representative donor in static (Static) and perfused chips (Flow) after 4 days of culture (similar results obtained in 3 donors).

**Supplementary Figure 4.**
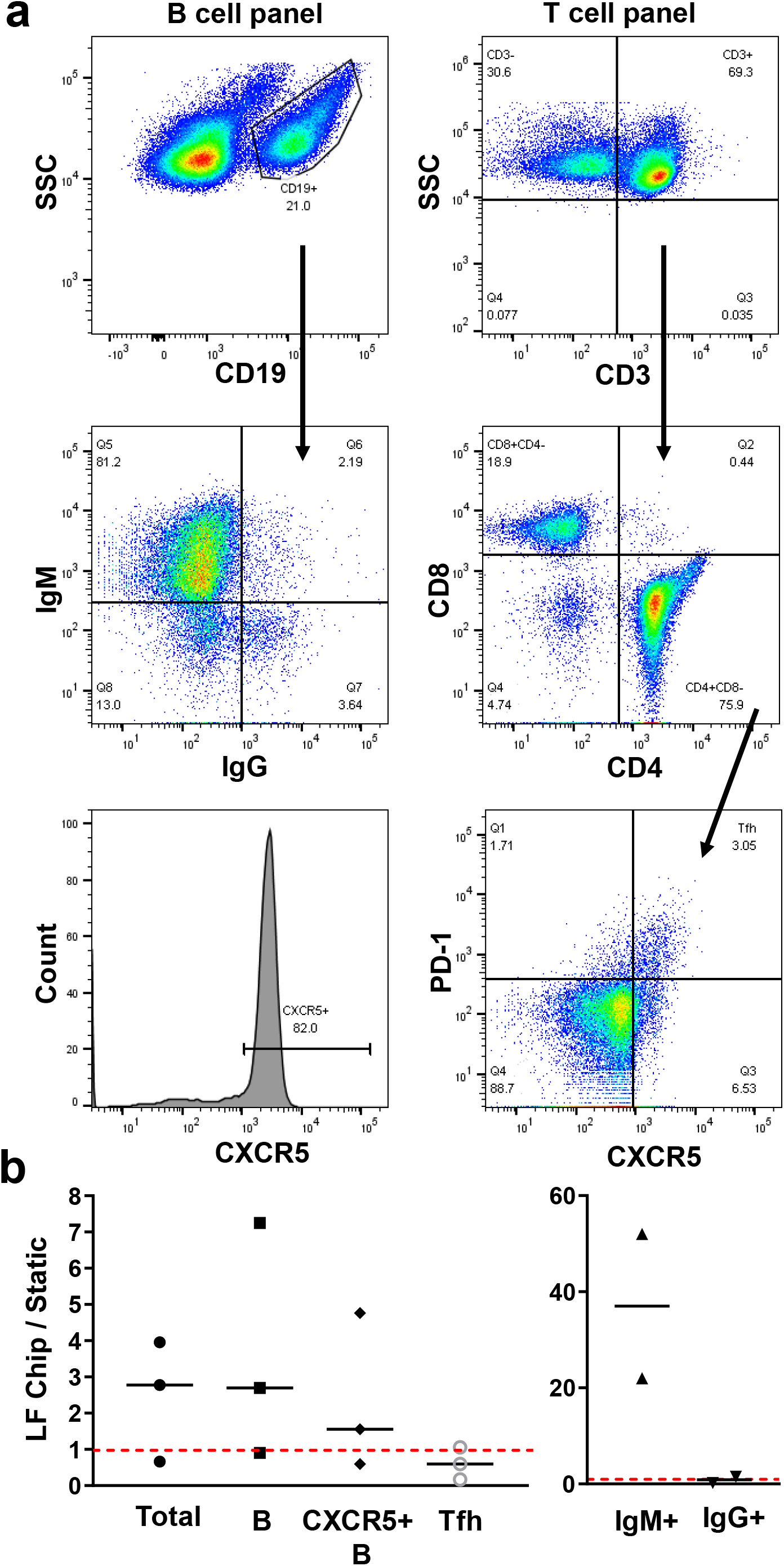
Flow cytometric analyses of the human LF chip. **a)** Dot plots of chips stained with a viability dye and fluorophore labeled antibodies against CD19, CD3, CD4, CD8, CXCR5, IgM, IgG and PD-1. **b)** Relative number of cells found on the LF chip compared to 2D culture. Each data points represents the ratio of the number of cells found in one chip vs. 2D well for an independent donor.

**Supplementary Figure 5.**
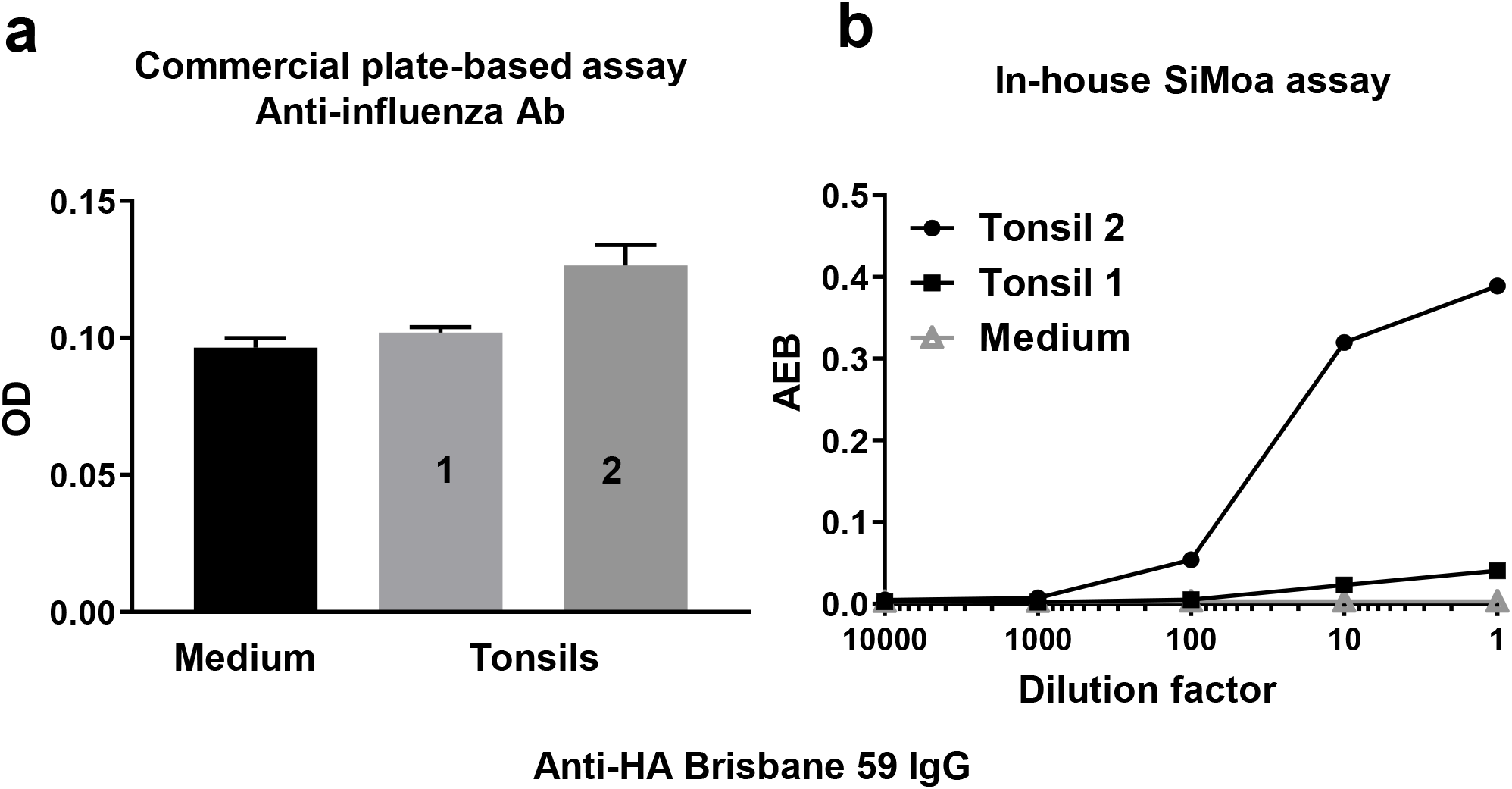
Comparison of anti-influenza antibody ELISAs. Results from tonsil supernatants (2 donors) and medium control shown. **a)** Optical density of samples (OD) in a commercially available ELISA for antibodies against influenza-A. **b)** Average Enzymes per Bead (AEB) obtained in a digital ELISA at different dilutions of the tonsil supernatant or medium control.

**Supplementary Figure 6.**
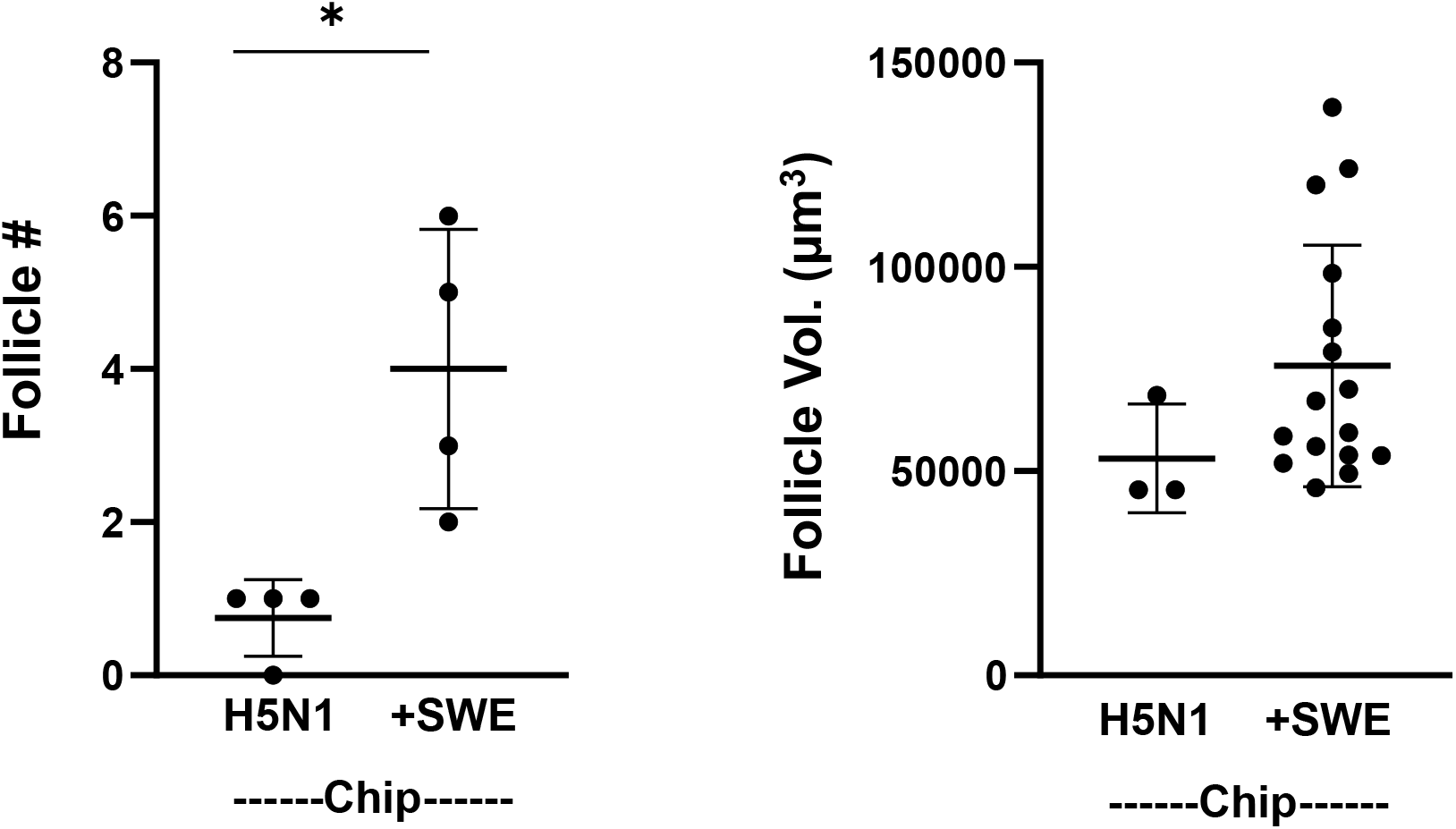
Effect of SWE on sustaining LFs. Follicle number (#, left) from 2 independent fields of view each from 2 LF chips from a donor vaccinated with split virion H5N1 alone or with 10 μl/ml SWE. Follicle Volume (Vol., right) was measured; each dot represents one follicle.

